# Towards a foundation model for raw nanopore signal through discrete tokenization

**DOI:** 10.64898/2026.11.05.764536

**Authors:** Guangzhao Cheng, Esa Pitkänen

## Abstract

Nanopore sequencing reads nucleic acids as changes in ionic current that carry information beyond the base sequence. Self-supervised pretrained models have advanced genomic sequence modeling and single-cell analysis, but remain underexplored for raw nanopore signals, which are continuous and lack a native token vocabulary. Here we present Brieno, an approach combining reconstruction-based tokenization through residual vector quantization with a Transformer for contextual representation learning. We pretrain the Transformer using masked-token prediction on raw nanopore signals from 118 million reads. With frozen or fine-tuned features, the model identifies species without requiring reference genomes. It complements Dorado by accurately classifying reads left unclassified in native-barcode demultiplexing, approaches Dorado’s basecalling performance across six genomes, and exceeds Dorado’s specialized 5-methylcytosine model as well as a dedicated 6-methyladenine caller. Brieno also supports accurate CpG methylation detection with limited labeled data. Across tasks, pretrained contextual representations improve downstream performance over non-pretrained baselines and generalize across sequencing runs. Together, these findings support self-supervised pretraining as a basis for transferable representations of raw nanopore signals.

## 1. Introduction

Nanopore sequencing, exemplified by Oxford Nanopore Technologies (ONT), measures changes in ionic current as a DNA or RNA strand passes through a protein pore, generating a continuous signal from which the nucleotide sequence can be inferred [1–3]. Beyond the base sequence, nanopore signals also reflect nucleotide modifications [4, 5], nucleic acid structure [6] and translocation dynamics [7]. Analyses of these signals support diverse applications, including species identification [8, 9] and RNA structure probing [10]. However, existing approaches typically rely on task-specific models or reference-based methods. For example, modification callers identify modified bases from characteristic signal patterns [4, 5, 11], demultiplexers recognize barcode signals at read ends [12, 13], and species classifiers match signals to references [8, 9, 14, 15] or learn from labeled reads [16, 17]. Despite operating on the same type of raw nanopore signal, these approaches are typically developed for individual tasks, with limited reuse of learned representations across tasks.

One promising approach to address this challenge is self-supervised pretraining, which enables foundation models to learn representations from large datasets and transfer them across tasks. This approach has advanced representation learning across text [18], speech [19, 20], genomic [21] and protein sequences [22], and single-cell transcriptomes [23, 24]. Speech offers a useful analogy to nanopore signals: both are continuous measurements that encode an underlying discrete sequence. Advances in speech representation learning [19, 25] and neural audio codecs [26, 27] suggest that discretization can support contextual representation learning from continuous signals. In the nanopore field, NanoBaseLib [28] and RawBench [29] already provide datasets and evaluation frameworks to support raw signal model development. However, extending self-supervised pretraining to nanopore signals presents distinct challenges. Raw signals lack a native token vocabulary, and sequence-dependent variation is intertwined with effects of nucleotide modifications, translocation dynamics and measurement noise. Discretization must therefore preserve information relevant to multiple tasks, and the benefits of pretraining must be established through downstream evaluation [30, 31].

Here we present Brieno, a self-supervised model for learning transferable representations from raw nanopore DNA signals. Brieno combines a residual vector-quantized tokenizer trained through signal reconstruction with a Transformer pretrained using masked-token prediction on signals from 118 million reads (Fig. 1a). We characterize how tokenization compresses signals while preserving sequence information, and evaluate frozen and fine-tuned representations on species classification, native-barcode demultiplexing, basecalling and DNA methylation detection. Comparisons with established methods on held-out datasets assess performance across sequencing runs, devices and biological samples. We further examine how quantization, contextual representation learning and fine-tuning affect downstream performance, and evaluate the utility of pretrained representations when labeled data are limited. Our results show that Brieno learns representations that transfer across diverse signal analysis tasks and support accurate CpG methylation detection with limited labeled data. These findings highlight the potential of discrete tokenization and self-supervised pretraining for developing foundation models of raw nanopore signals.

**Figure 1:**
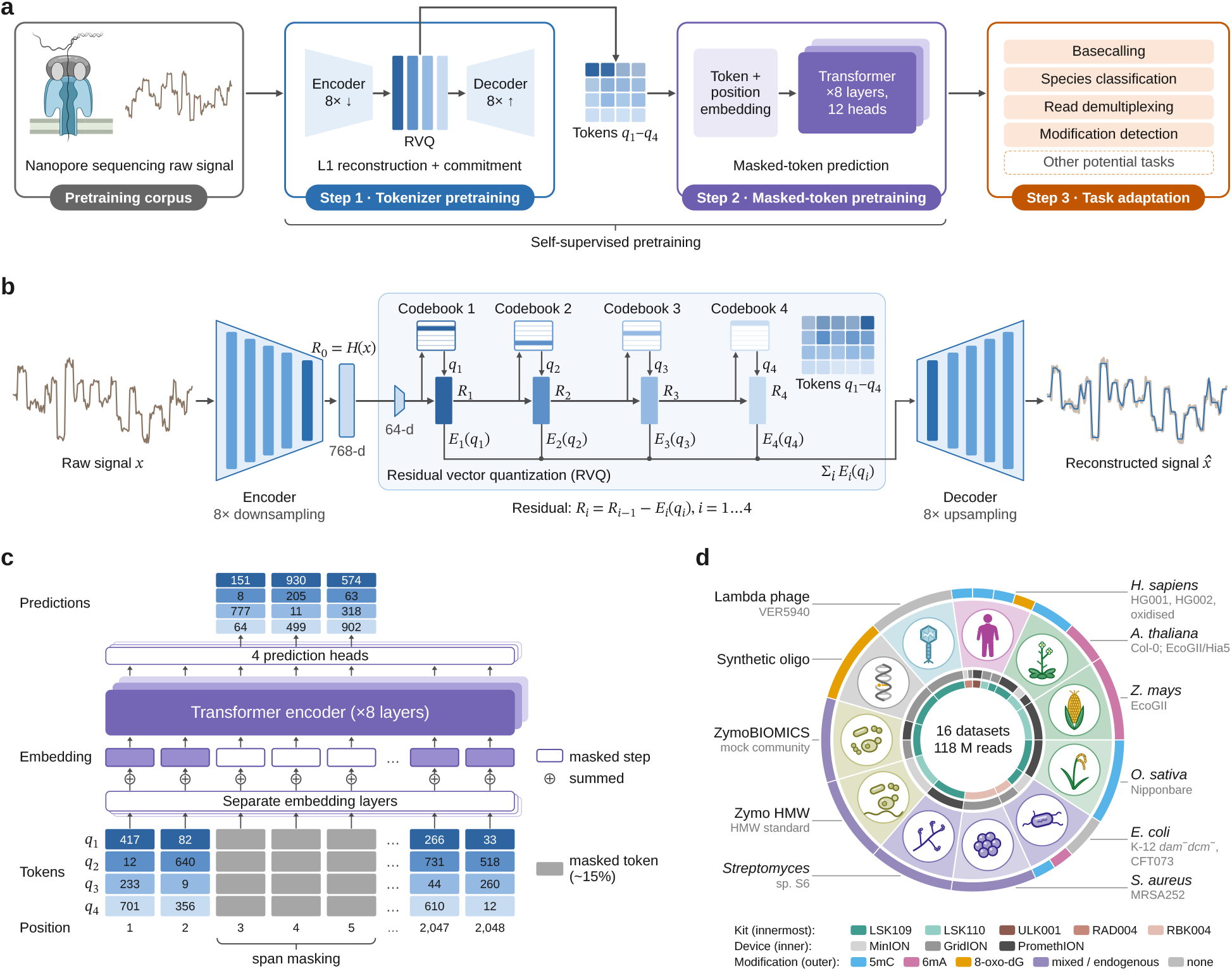
Overview of Brieno. **a**, The workflow of Brieno. Self-supervised pretraining proceeds in two stages: a tokenizer learns to convert raw nanopore signals into discrete tokens, and a Transformer learns contextual representations through masked-token prediction. The pretrained model is subsequently used as a frozen feature extractor or fine-tuned for downstream tasks, including basecalling, species classification, read demultiplexing and modification detection. **b**, Tokenizer architecture. The encoder output is quantized by four residual vector quantization (RVQ) stages, each selecting a code *q*_*i*_ from its own codebook, and the decoder reconstructs the signal from the summed code vectors. **c**, Masked-token pretraining. Spans of positions are masked across all four codebooks, per-codebook embeddings are summed at each position (*⊕* ) and a Transformer with four prediction heads predicts the masked tokens. **d**, Diagram of the pretraining corpus, 16 nanopore sequencing datasets (118 million reads), annotated by source organism, library kit, sequencing device and DNA modification. Supplementary Table S1 lists all datasets used in this study.

## 2. Results

### 2.1. Brieno model architecture and pretraining

Brieno combines signal tokenization with masked-token pretraining to learn representations of raw nanopore signals in two self-supervised steps (Fig. 1a). In the first step, a tokenizer inspired by neural audio codecs [26, 27] converts raw nanopore signals into discrete token sequences. A convolutional encoder downsamples the raw current signal eightfold into a sequence of latent frames, with one feature vector per eight input samples. A residual vector quantizer [32] assigns four tokens to each frame, one from each of four 1,024-entry codebooks. A decoder reconstructs the signal from the summed code vectors. The tokenizer is trained with an *L*_1_ reconstruction loss, the mean absolute reconstruction error, and a commitment loss, which encourages encoder outputs to remain close to their quantized representations (Fig. 1b; Methods).

In the second step, a Transformer with eight layers, 12 attention heads per layer and 61.4 million parameters is pretrained using masked-token prediction [18]. Contiguous spans are masked across all four codebooks, and per-codebook embeddings are summed at each position. Four prediction heads predict the masked tokens from the Transformer outputs, encouraging the model to learn contextual representations (Fig. 1c). For downstream applications, a task-specific head is trained either on frozen Transformer features or jointly with the Transformer through fine-tuning, while the tokenizer remains frozen in both settings.

For self-supervised training and validation, we assembled 16 public datasets comprising 118 million reads from ONT R9.4.1 flow cells. The collection spans bacteria, fungi, plants, humans, synthetic oligonucleotides and two mock communities, across three sequencing platforms and five library preparation kits, and includes 5mC, 6mA and 8-oxo-dG modifications (Fig. 1d, Supplementary Table S1). Twelve datasets were used for training and four were held out for validation. Neither self-supervised step requires basecalls, alignments or modification labels.

### 2.2. Discrete tokenization preserves sequence information in nanopore signals

To assess how much signal information is preserved during tokenization, we first evaluated reconstruction quality on held-out validation data. The tokenizer achieved a reconstruction *L*_1_ error of 0.030 in normalized current units, with reconstructed signals closely following the raw trace (Fig. 2a). Reconstruction progres-sively improved as additional residual codebooks were included: for the illustrated window, the *L*_1_ error decreased from 0.107 using the first codebook alone to 0.030 using all four (Supplementary Fig. S1a). The first codebook captured the main current levels, whereas subsequent codebooks refined the reconstruction.

**Figure 2:**
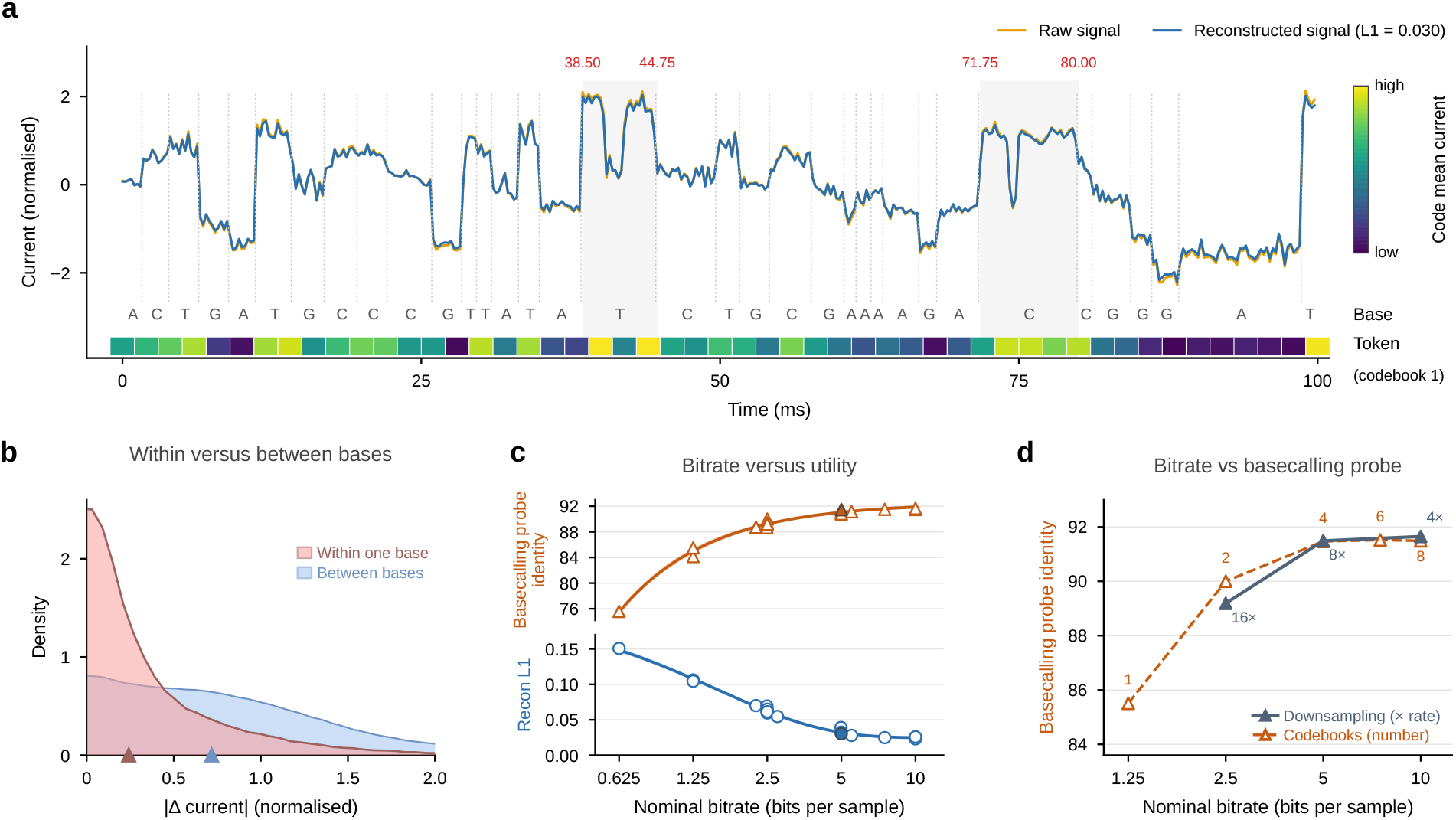
Design of the Brieno tokenizer. **a**, Comparison of raw (orange) and reconstructed (blue) signals for a 100 ms window from a held-out bacteriophage lambda read. Dotted lines mark reference-position boundaries determined by Uncalled4. Reference bases, taken as the third base of the Uncalled4 6-mer, and first codebook tokens are shown below, with tokens colored by the mean current associated with each code. Shaded regions highlight current variation within individual reference positions, in which red labels indicate their start and end times. **b**, Distributions of absolute current differences within one base (red) and between consecutive bases (blue) across four species. Triangles indicate medians. **c**, Reconstruction error (blue circles) and basecalling probe identity (orange triangles) versus nominal bit rate for 21 tokenizers. Filled markers indicate the selected tokenizer. Curves show three-parameter nonlinear least-squares fits. **d**, Effects of encoder downsampling and codebook number on basecalling probe identity, on the same nominal bit-rate axis. Downsampling is varied with four codebooks (solid line), and codebook number is varied at 8 *×* downsampling (dashed line). These comparisons guided the choice of 8 *×* downsampling rate and four codebooks.

We next examined how tokens relate to the underlying nucleotide sequence. We aligned 8,000 held-out signal chunks from four species to their references using Uncalled4 [7], which maps each aligned signal segment to a reference 6-mer. We labeled each reference position by the third base of its 6-mer, as this position explained the largest fraction of current variance (Supplementary Fig. S1b). Across 9.75 million reference positions, each position was covered by an average of 1.68 latent frames, and 41% were covered by multiple frames. We focused on the first codebook token assigned to each frame. The two shaded regions (38.50–44.75 ms and 71.75–80.00 ms) in Fig. 2a illustrate how multiple frames and their associated tokens span a single reference position with varying current levels. Among positions covered by exactly two frames, the frames received different tokens in 99.5% of cases. Representing each token by its code-associated mean current, we observed median absolute current differences of 0.24 within reference positions and 0.72 between consecutive positions (Fig. 2b; Methods). The corresponding median absolute differences calculated from frame-level raw currents were 0.23 and 0.69 respectively, indicating that tokenization largely preserved the median magnitude of both within-position and between-position variation. Thus, first codebook tokens retain current variation within individual reference positions rather than serving as one-to-one labels for bases.

To assess whether improved signal reconstruction translates into better sequence decoding, we compared reconstruction error and basecalling probe identity, measured with a small decoder trained on the frozen tokens (see Methods), across 21 tokenizer configurations (Fig. 2c and Supplementary Table S2). We define nominal bit rate as the number of codebooks multiplied by the log_2_ codebook size and divided by the downsampling factor. Relative to the 16-bit representation of raw current samples, 5 bits per sample corresponds to a nominal 3.2-fold compression. Across approximately 0.6–10 bits per sample, reconstruction error generally decreased with increasing bit rate, whereas basecalling probe identity largely plateaued above approximately 5 bits per sample. Thus, additional capacity improved signal reconstruction without consistently improving sequence decoding, motivating tokenizer selection based on both reconstruction quality and downstream utility.

We next examined how encoder downsampling and codebook number affected basecalling probe identity (Fig. 2d). At a matched bit rate of 2.5 bits per sample, eightfold downsampling with two codebooks achieved 90.00% identity, compared with 89.19% for sixteenfold downsampling with four codebooks, indicating that temporal resolution also influenced sequence decoding. At eightfold downsampling, increasing the number of codebooks from four to six or eight yielded gains of only 0.05 and 0.02 percentage points, respectively. Reducing downsampling to fourfold provided a modest additional gain but doubled the sequence length supplied to the Transformer. We therefore selected eightfold downsampling with four 1,024-entry codebooks, corresponding to 5 bits per sample.

### 2.3. Brieno enables species classification and barcode demultiplexing without basecalling

We first evaluated whether Brieno representations support species classification directly from raw signals without reference genomes at inference. Using the ZymoBIOMICS mock communities [33] (D6300, ten species; D6322, eight of the same species), we trained a species classification head on five sequencing runs, either with frozen Brieno representations or jointly with the Transformer through fine-tuning. We evaluated both settings on two held-out runs: a Quadram GridION run of D6300 containing 3.73 million reads and a Johns Hopkins MinION run of D6322 containing 634,375 reads, which carries eight of the ten species while the classifier retained all ten outputs. All methods were evaluated on the same read sets, with unclassified reads counted as errors. Fine-tuned Brieno achieved read-level accuracies of 0.9887 and 0.9965, while Brieno with frozen representations achieved 0.9728 and 0.9916, respectively (Fig. 3a). Both exceeded RawHash2 [9], the strongest reference-based comparator, which achieved 0.9564 and 0.9891. These results demonstrate that Brieno representations support accurate species classification across held-out sequencing runs without signal-to-reference mapping.

**Figure 3:**
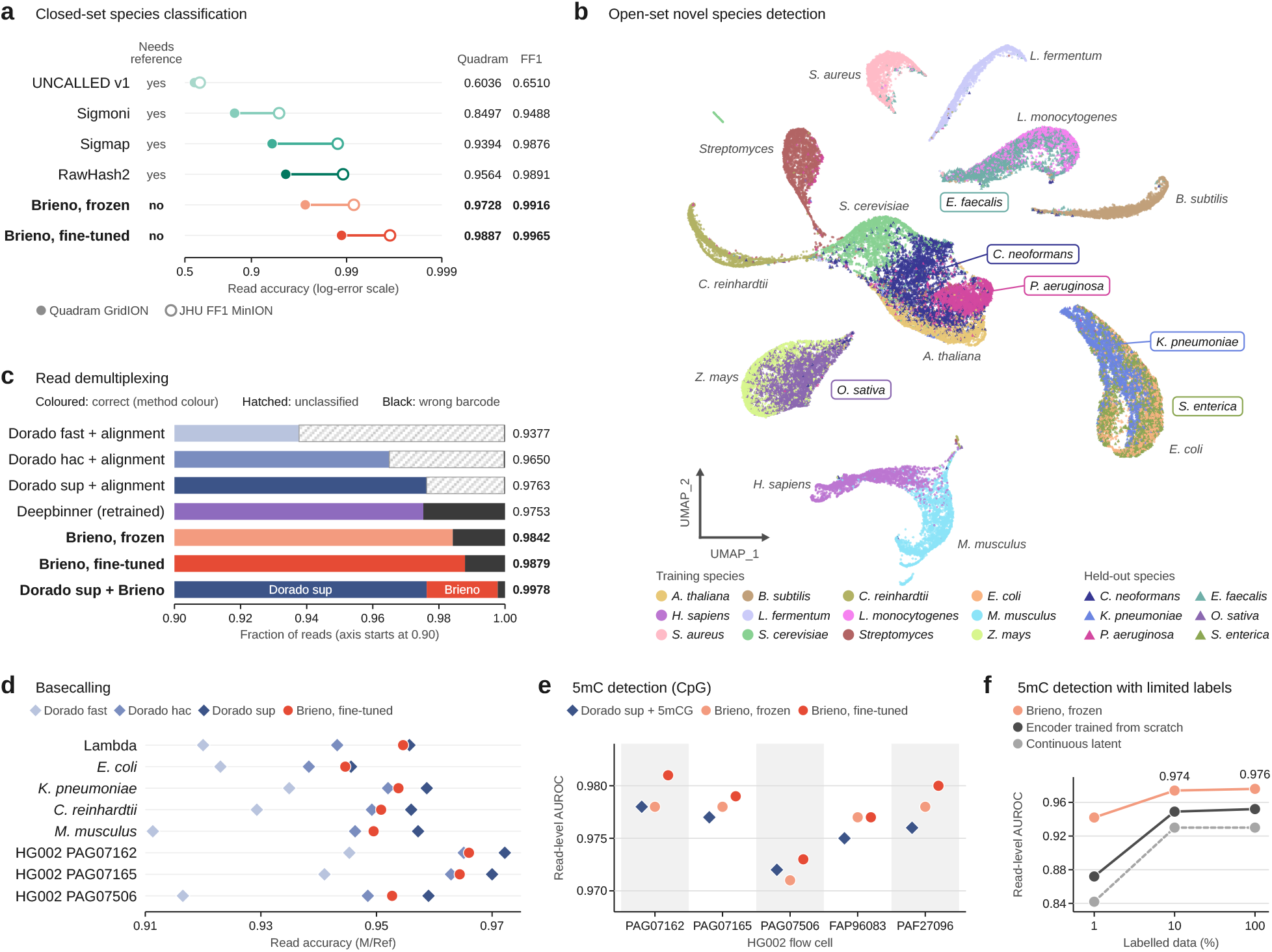
Evaluation of Brieno across downstream tasks. **a**, Closed-set species classification: read-level accuracy on two held-out sequencing runs, Quadram GridION (filled circles) and JHU FF1 MinION (open circles). Reference-genome requirements are indicated for each method. **b**, Open-set novel species detection: UMAP projection of read embeddings for twelve training classes and six held-out species. Colors indicate species identity; circles denote training classes and triangles denote held-out species, whose labels are boxed. **c**, Native-barcode demultiplexing on two held-out flow cells. Stacked bars show the fractions of correctly classified, unclassified and incorrectly classified reads; numbers indicate overall accuracy. The bottom bar is a cascade in which Dorado at super accuracy keeps the reads it classified and fine-tuned Brieno classifies the reads it refused. The horizontal axis starts at 0.90. **d**, Read-level basecalling accuracy across species and human HG002 sequencing runs, comparing fine-tuned Brieno with Dorado fast, hac and sup models. **e**, Read-level AUROC for 5mC detection at CpG sites across five held-out HG002 flow cells, comparing frozen and fine-tuned Brieno with Dorado. **f**, Read-level CpG AUROC on HG002 as a function of the fraction of HG001 training chunks retained, at a fixed optimization budget, for frozen Brieno, an encoder trained from scratch and the continuous latent.

We next examined whether these representations support the detection of species absent from classifier training. We trained a separate twelve-class model on sequencing runs of individual species and of the ZymoBIOMICS mock communities, spanning mammals, plants, algae, fungi and bacteria. Fine-tuned Brieno achieved a read-level accuracy of 0.9803 on held-out sequencing runs, while Brieno with frozen representations achieved 0.9776. We then examined representations of six additional species excluded from classifier training. The UMAP projection revealed species-associated structure among these held-out reads (Fig. 3b). *O. sativa* reads were enriched in a localized region near *Z. mays*, while *P. aeruginosa* reads formed a concentrated region within the broader embedding. However, separation varied across species, with *S. enterica* reads extensively overlapping those of *E. coli*. To detect unseen species, we used the negative margin between the two highest read-level class scores as a novelty score, with smaller margins indicating greater uncertainty. This score distinguished unseen from known species with an overall AUROC of 0.72, ranging from 0.54 for *S. enterica* to 0.88 for *C. neoformans* (Supplementary Fig. S2a and Supplementary Methods). These results indicate that the representations capture species-associated structure beyond the training classes, although their utility for detecting unseen species varies across species.

We also evaluated native-barcode demultiplexing using 24 ONT barcodes across eight flow cells. Six flow cells were used for training and two for testing, with disjoint isolate sets across flow cells to reduce the possibility of predicting barcodes from isolate-specific genomic signals. All methods were evaluated on the same 1,998,079 test reads without quality or length filtering. Fine-tuned Brieno classified every read and achieved an overall accuracy of 0.9879, while Brieno with frozen representations achieved 0.9842. Both exceeded Dorado super-accuracy model followed by barcode alignment [3] (0.9763) and a retrained Deepbinner model [12] (0.9753; Fig. 3c). These comparisons count an unclassified read as an error, and the two routes make different kinds of errors: Dorado at super accuracy left 2.35% of reads unclassified and misassigned 0.02%, whereas fine-tuned Brieno classified every read and misassigned 1.21% (Fig. 3c). When Brieno was allowed to abstain and its coverage was matched to the 97.65% of reads Dorado classified, Dorado remained the more precise of the two on the reads it called (0.9997 against 0.9982); the advantage of Brieno is therefore one of coverage rather than of per-read precision. Brieno correctly assigned 91.7% of the 46,910 reads left unclassified by Dorado at super accuracy. A cascade that retained Dorado assignments and used fine-tuned Brieno only for unclassified reads achieved an overall accuracy of 0.9978 (Fig. 3c). These results demonstrate that Brieno supports accurate native-barcode demultiplexing and complements sequence-based pipelines by assigning reads they leave unclassified.

### 2.4. Brieno adapts pretrained signal representations for basecalling

We next evaluated whether Brieno representations support nucleotide sequence decoding. We attached a CTC-CRF decoder [34] to the pretrained Transformer and trained them jointly on four datasets, while keeping the tokenizer frozen. Training labels were obtained through signal-to-reference alignment using Uncalled4 [7]. We compared Brieno with Dorado (v0.9.6) at fast, high-accuracy (hac) and super-accuracy (sup) settings [3] on identical read sets using two evaluation protocols with three scoring settings. For per-read evaluation, each basecalled read was aligned to a corresponding reference sequence extracted from the genome using Dorado sup alignments. We calculated accuracy as matched bases divided by either the reference sequence length, used as the primary metric (Fig. 3d and Supplementary Table S3), or the alignment length, which additionally accounts for insertions (Supplementary Table S4). The third setting scores alignments to the whole genome (Supplementary Table S5; Supplementary Methods).

Fine-tuned Brieno approached Dorado sup’s basecalling accuracy under the primary metric, achieving 0.9546 versus 0.9557 on bacteriophage lambda and 0.9446 versus 0.9456 on *E. coli* (Supplementary Table S3). The gaps were larger on *K. pneumoniae* and *C. reinhardtii*, at 0.0049 and 0.0052, and ranged from 0.0056 to 0.0077 on the three HG002 flow cells and *M. musculus*. Nevertheless, Brieno exceeded Dorado hac across all eight evaluation groups under each of the three scoring settings (Supplementary Tables S3–S5). These results demonstrate that Brieno’s pretrained representations support accurate basecalling across diverse species and sequencing runs, approaching the accuracy of a specialized basecaller while retaining applicability to other signal analysis tasks.

### 2.5. Brieno supports accurate DNA methylation detection with limited labeled data

We next evaluated whether Brieno representations support DNA methylation detection, beginning with 5-methylcytosine (5mC) at CpG sites. We trained a modification detection head on HG001 reads using pseudo-labels derived from whole-genome bisulfite sequencing, either with frozen Brieno representations or jointly with the Transformer through fine-tuning. The model predicts one methylation score per CpG site per read. Evaluation included 1.83 million read–site pairs from five HG002 flow cells excluded from all training stages, using EpiQC high-confidence methylation labels [35]. The test set comprised four PromethION flow cells and one GridION flow cell prepared with a different library kit. Fine-tuned Brieno achieved an average read-level AUROC of 0.978, compared with 0.976 for frozen representations and 0.975 for Dorado’s dedicated 5mCG model. It exceeded Dorado on all five flow cells (Fig. 3e), supporting transfer across sequencing runs. We evaluated individual read–site predictions because the sampled test reads provided insufficient coverage for reliable site-level methylation frequency estimation (Methods).

We then assessed how detection performance changed when labeled training data were limited. Using 10% or 1% of the HG001 training chunks while keeping the fine-tuning step budget fixed, frozen Brieno representations achieved pooled HG002 AUROCs of 0.974 and 0.942, respectively (Fig. 3f). With 10% of the labels, performance remained close to the full-data AUROC of 0.976. With 1%, corresponding to approximately 29,000 labeled chunks, frozen Brieno exceeded an encoder trained from scratch by 0.070 AUROC (0.942 versus 0.872). Its performance was also only 0.010 below that of the encoder trained from scratch using the full set of approximately 2.9 million labeled chunks. These results demonstrate that frozen pretrained representations support accurate CpG methylation detection with substantially fewer labeled examples.

We further evaluated 6-methyladenine (6mA) detection at GATC sites using the NEMO datasets [11]. Models were trained on *E. coli* GridION datasets and tested on *E. coli* PromethION and *H. pylori* datasets. Fine-tuned Brieno achieved read-level AUROCs of 0.991 and 0.990, respectively, while frozen representations achieved 0.985 and 0.986. Both exceeded the NEMO GATC model, which achieved 0.947 and 0.917 on the same evaluation datasets (Supplementary Fig. S2b). Performance varied across other methylated motifs in *H. pylori*, suggesting that transfer beyond the GATC training context was limited. Together, these results support the application of Brieno representations to both cytosine and adenine methylation detection, with cross-species transfer demonstrated within the GATC sequence context.

### 2.6. Pretrained contextual representations improve downstream performance

To examine how quantization, contextual representation learning and pretraining contribute to downstream performance, we compared features from different stages of Brieno using the same task-specific head architecture (Fig. 4a; Supplementary Methods). These comprised the continuous encoder latent before quantization (A), the quantized latent (B1), discrete tokens with learned embeddings (B2), frozen pretrained Transformer features (C1) and fine-tuned Transformer features (C2). We included two controls: a randomly initialized Transformer trained with the task head while keeping the tokenizer frozen (D1), and an encoder trained from scratch with the task head (D2). These comparisons assess how the representation supplied to the head affects performance and distinguish the benefits of Transformer pretraining from those of the architecture itself.

**Figure 4:**
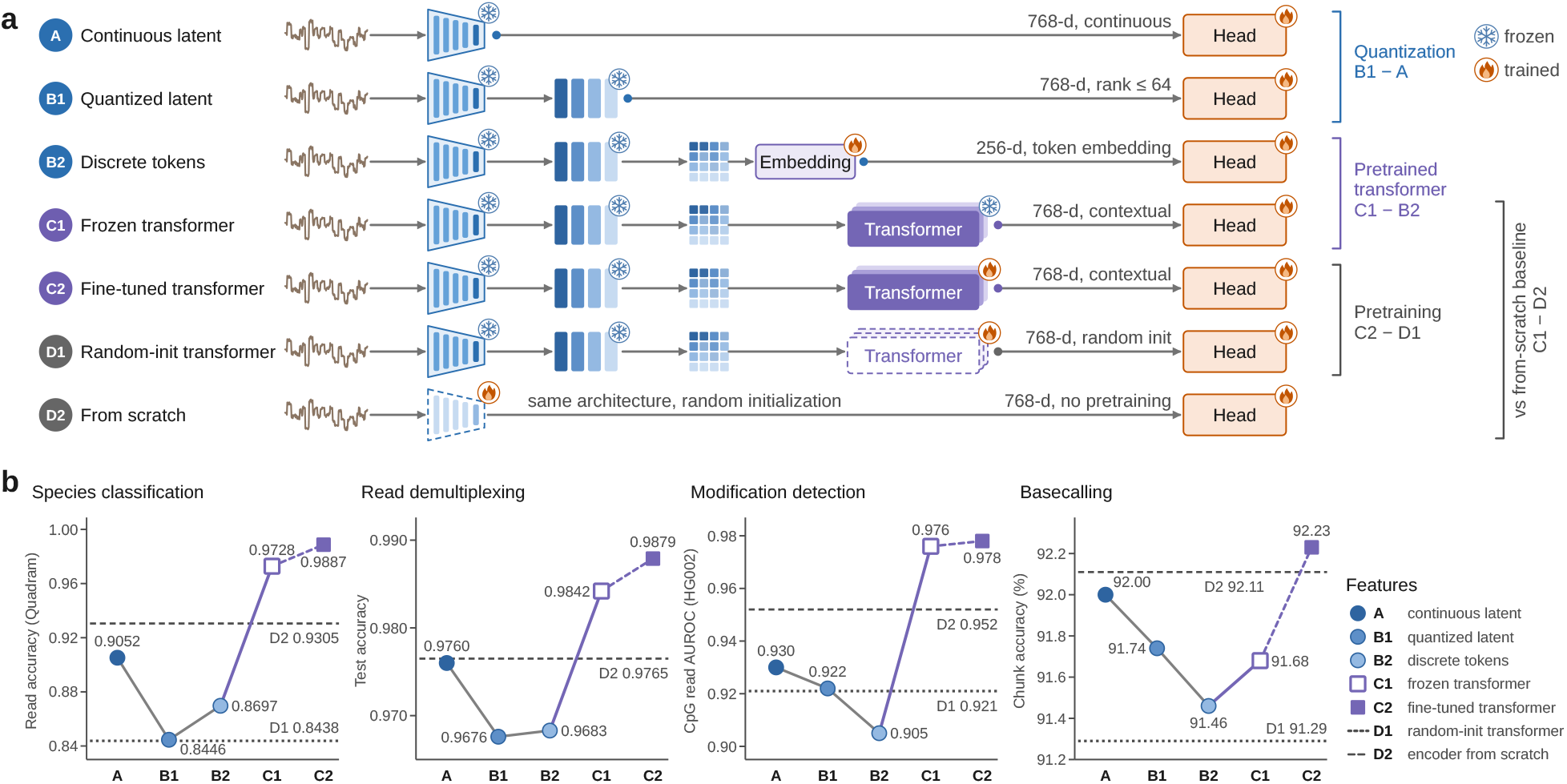
Effects of quantization and pretraining on downstream performance. **a**, Feature representations and training configurations used for comparison: continuous encoder latents (A), quantized latents (B1), discrete tokens with learned embeddings (B2), frozen Transformer features (C1), fine-tuned Transformer features (C2), a randomly initialized Transformer trained with the head (D1) and an encoder trained from scratch (D2). Each representation is evaluated using the same task-specific head architecture. Snowflakes and flames indicate frozen and trainable components, respectively. **b**, Performance on species classification, read demultiplexing, DNA modification detection and basecalling using different features. Dashed lines indicate the randomly initialized Transformer (D1) and the encoder trained from scratch (D2).

Quantization reduced performance across all four tasks, although the magnitude varied (Fig. 4b). Relative to the continuous latent, the quantized latent reduced species classification accuracy by 6.06 percentage points, demultiplexing accuracy by 0.84 percentage points, methylation detection AUROC by 0.008 and basecalling accuracy by 0.26 percentage points. Frozen pretrained Transformer features nevertheless out-performed the discrete tokens on every task, and the continuous latent on all but basecalling. Compared with discrete tokens, these features improved species classification accuracy by 10.31 percentage points, demultiplexing accuracy by 1.59 percentage points, methylation detection AUROC by 0.071 and basecalling accuracy by 0.22 percentage points. Fine-tuning further improved performance across all four tasks. Thus, frozen pretrained features more than offset the performance loss from quantization on three of the four tasks, and fine-tuned features did so on all four.

To assess the contribution of pretraining while controlling for Transformer architecture, we compared fine-tuned Brieno with the randomly initialized Transformer. For methylation detection, the pretrained model achieved an AUROC of 0.978, compared with 0.921 for the randomly initialized control under the same training data and optimization step budget, a gain of 0.057 (Fig. 4b). Frozen pretrained features also outperformed the encoder trained from scratch, which achieved an AUROC of 0.952. Doubling the training budget increased the latter to 0.964, narrowing the gap but remaining below the 0.976 achieved with frozen pretrained features. These results indicate that training budget explains part of the difference from the encoder baseline, while the architecture-matched comparison provides evidence for the contribution of self-supervised pretraining to methylation detection. On read demultiplexing the randomly initialized

Transformer remained at the 24-class chance level across a tenfold range of learning rates, as did the discrete-token rung when its embedding was initialized randomly rather than from the tokenizer’s codebooks, whereas the encoder trained from scratch on the raw signal reached 0.9765. These results suggest that an informed token-embedding initialization facilitates optimization on this task under the training settings examined.

## 3. Discussion

We developed Brieno, a self-supervised framework that combines discrete signal tokenization with Transformer pretraining to learn reusable representations of raw nanopore signals. A shared pretrained backbone supported four downstream applications through task-specific heads. Frozen representations sufficed for all four tasks, and fine-tuning further improved performance on each of them. Pretrained models outperformed randomly initialized Transformers with identical architecture, confirming the benefit of self-supervised pre-training. Beyond individual benchmarks, Brieno complemented sequence-based demultiplexing by assigning reads left unclassified by Dorado and supported accurate CpG methylation detection with limited labeled data. Our analyses also characterized the trade-offs between signal reconstruction, nominal bit rate and downstream utility, informing tokenizer design and pretraining for raw nanopore signals.

An alternative to discrete tokenization is to supply continuous encoder features directly to the Transformer. In Brieno, discretization provides a finite vocabulary for masked-token prediction but introduces information loss. Frozen Transformer representations outperformed continuous encoder features on three of the four tasks. For basecalling, only the fine-tuned representations surpassed them (Fig. 4b). These comparisons do not establish that discrete inputs are preferable to continuous inputs for Transformer pretraining. Following approaches in speech [19, 20], continuous features could instead serve as inputs while discrete representations provide prediction targets; we left evaluation of this setting under matched architectures and pretraining data as future work. Discrete inputs nevertheless offer possibilities beyond the predictive tasks evaluated here. A finite vocabulary enables token-based sequence modeling, potentially supporting conditional signal simulation given a nucleotide sequence, modification state or sequencing condition. A fixed codebook could also provide a shared representational reference for adapting encoders across new pore chemistries.

We pretrained Brieno on R9.4.1 signals to take advantage of the extensive publicly available datasets. To explore its potential as a general-purpose foundation model applicable across pore chemistries, we conducted an initial evaluation on R10.4.1 data. This evaluation suggests that the pretrained representations retain utility for species classification after supervised adaptation (Supplementary Fig. S3). Future work could explore chemistry-specific encoders, continued self-supervised pretraining and joint pretraining across chemistries to broaden the applicability of Brieno.

## 4 Methods

### Pretraining corpus and signal preprocessing

The pretraining corpus comprises 16 public nanopore datasets generated on ONT R9.4.1 flow cells, spanning bacteria, fungi, plants, humans, synthetic oligonu-cleotides and two mock communities, across three sequencing platforms and five library preparation kits, and including 5mC, 6mA and 8-oxo-dG modifications. Twelve were used for training and four held out for validation. Each read is normalized by subtracting its median and dividing by its median absolute deviation, then divided into chunks of 16,384 samples with an overlap of 2,048 samples. Reads shorter than one chunk are discarded. More details are listed in Supplementary Table S1.

### Tokenizer design and evaluation

The tokenizer follows EnCodec [27]. Here *T* denotes the number of samples in a chunk, *d* the encoder downsampling factor, *n*_*q*_ the number of residual codebooks, *K* the number of entries per codebook and *d*_*q*_ the dimension in which quantization is performed. A convolutional encoder with log_2_ *d* stride-2 blocks and a final stride-1 projection produces *N* = *T/d* latent frames of width 768 from a chunk *x∈* ℝ^*T*^ . These frames are then projected to *d*_*q*_ dimensions for residual vector quantization, giving *z*. A residual vector quantizer [26, 32] with *n*_*q*_ codebooks 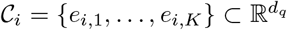 quantizes every frame *z*_*n*_ by successive nearest-neighbor search on the residual,

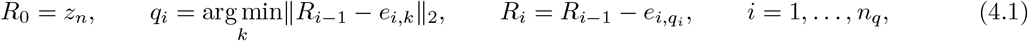

so the frame becomes the token tuple (*q*_1_, …, *q*_*n*_ ) and is reconstructed as 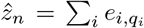 . Codebooks are initialized with *k*-means and updated using an exponential moving average. The quantized frames are projected back to 768 dimensions, and a mirrored decoder reconstructs the chunk as 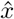. The projections into and out of the quantization space are optional: when *d*_*q*_ = 768 they are omitted and quantization acts directly on the encoder output, whereas the selected tokenizer sets *d*_*q*_ = 64, reducing its four codebooks from 3.1 M to 0.26 M parameters. The total loss for tokenizer pretraining is

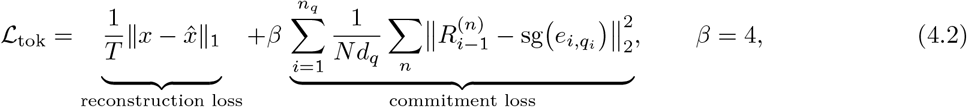

Where 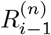is the residual of frame *n* entering quantizer *i*, sg denotes the stop-gradient operator, and a straight-through estimator passes the decoder gradient to the encoder. With *n*_*q*_ codebooks of *K* entries and one latent frame every *d* samples, each frame is described by *n*_*q*_ log_2_ *K* bits, so the nominal bit rate is

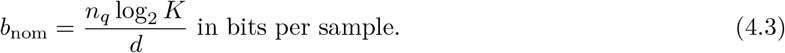

The 21 tokenizers of Fig. 2c,d vary *d ∈ {*4, 8, 16 }, *n*_*q*_ *∈ {*1, 2, 4, 6, 8}, *K∈ {* 512, 1,000, 1,024, 2,048 }and *d*_*q*_ *∈ {*32, 64, 768} at an encoder width of 768, except for one run that halves it to 384 (Supplementary Table S2). Token-level analyses (Fig. 2b and Supplementary Fig. S1) use 8,000 chunks from four species aligned to their references at the signal level with Uncalled4 [7]. Tokenizers were ranked by the sequence identity achieved by a basecalling probe trained on their frozen tokens and evaluated on 20,000 held-out lambda chunks (Supplementary Methods).

### Transformer pretraining

The Transformer is an 8-layer pre-LayerNorm encoder with 12 attention heads, a hidden dimension of 768, a context length of 2,048 frames and 61.4 M parameters. The input at position *n* is 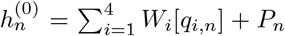, with *W ∈* ℝ ^(*K*+1)*×*768^ the embedding table of codebook *i*, whose extra row is that codebook’s learned mask embedding, and *P* a learned positional embedding. We mask a set of positions *ℳ* covering 15% of the sequence, using spans of at least 12 frames to discourage trivial interpolation from neighboring frames. Masked positions are corrupted as in BERT [18]: at 80% of them the token of every codebook is replaced by that codebook’s mask token, at 10% each codebook receives an independently sampled random token, and at 10% the tokens are left unchanged. The positional embedding is always retained, and the loss is computed at every masked position, including those left unchanged. The model is trained with the following masked-token prediction (MTP) loss:

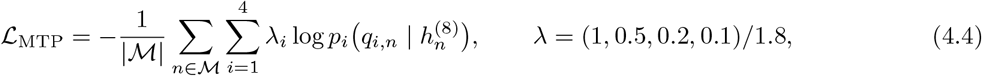

where 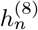 is the final-layer representation at position *n, p*_*i*_ is the softmax output of the *i*-th prediction head, and the weights *λ* downweight the later, residual codebooks. All downstream results use the epoch-70 checkpoint. Details of the prediction heads and the optimization settings are provided in Supplementary Methods.

### Species identification and barcode demultiplexing

The two tasks use separately trained heads with the same architecture: three strided convolutions, attention and mean pooling, and a two-layer classifier. Each read is assigned to the class with the highest mean log-probability across its chunks. For species identification, we used seven sequencing runs of the ZymoBIOMICS mock communities [33] (D6300, ten species; D6322, eight of the same species) generated in three laboratories. Five runs were used for training and two for testing (ERR3518146, Quadram GridION; SRR13127889, Johns Hopkins MinION). Per-read species labels were obtained by aligning Dorado super-accuracy basecalls to a ten-species pan-reference using minimap2. The embedding visualization in Fig. 3b was generated using a separate twelve-class model trained on single-species and mock-community runs, with six species excluded from classifier training. Novelty was assessed using the negative margin between the two highest read-level mean log-probabilities. For barcode demultiplexing, we used eight flow cells from ENA project PRJEB42440, comprising 8.2 million reads from 188 isolates with pairwise disjoint isolate sets across flow cells. Each isolate was assigned one of 24 ONT native barcodes. Six flow cells were used for training, and two were held out for testing (FAK63737, MinION; FAK47252, GridION). We compared Brieno with Dorado (v0.9.6) [3], using its built-in barcode aligner, and a retrained Deepbinner [12]. All methods were evaluated on all 1,998,079 deposited reads from the two test flow cells, without quality or length filtering. More details are provided in Supplementary Methods.

### Basecalling

The basecalling head consists of a linear projection to width *d*, a stride-2 transposed convolution that upsamples the representations from 2,048 to 4,096 frames per chunk, five bidirectional LSTM layers with *h* units and a globally normalized linear CRF over 5-mer states [34]. The main benchmarks use *h* = *d* = 512 (35 M parameters), trained for 120 epochs. Training data comprise 30.0 million chunks of *E. coli* (unmethylated), HG001, *Streptomyces* sp. S6 and *S. aureus*. Per-chunk 6-mer labels were obtained by aligning the signals to reference sequences using Uncalled4 [7]. Test data comprise the lambda run of Pagès-Gallego and de Ridder [36], six *E. coli* and five *K. pneumoniae* clinical-isolate runs, five *C. reinhardtii* runs, one *M. musculus* run and three HG002 PromethION flow cells. Reads were basecalled from overlapping 16,384-sample chunks and decoded by CRF beam search of width 32. All methods were evaluated using the nanopore-benchmark protocol [36]. Predicted sequences were aligned to per-read reference segments extracted from the genome using Dorado sup alignments. Accuracy was calculated as matched bases divided by reference segment length and averaged over successfully aligned reads. Uncalled or unalignable reads were excluded (median 0.26%, maximum 1.18% for Brieno; none for Dorado sup) and scored zero only in the separate genome-level evaluation. Restricting evaluation to reads aligned by Dorado sup may favor that model. Further details are provided in Supplementary Methods.

### Modification detection

The modification detection head produces one logit per frame. The logit for each base is obtained by averaging the logits over the frames spanned by its Uncalled4-aligned events [7] (Supplementary Methods). Predictions therefore correspond to individual cytosines or adenines on individual reads, as in Dorado [3], nanopolish [4] and Rockfish [5]. For 5mC detection, training data comprised 2.9 million chunks from 25 HG001 ultra-long sequencing runs, with pseudo-labels derived from ENCODE GM12878 whole-genome bisulfite sequencing. Chromosomes 20 and 21 were reserved for validation. Test data comprised five HG002 flow cells excluded from all training stages. Predictions were evaluated on 1.83 million read–site pairs using EpiQC high-confidence CpG labels [35]. Performance was measured by AUROC across individual read–site pairs. Site-level methylation frequencies were not evaluated because the sampled test reads provided insufficient per-site coverage for reliable estimation. Dorado v0.9.6 with its R9.4.1 5mCG model was evaluated on the same reads using the same scoring procedure. For 6mA detection, training data comprised *E. coli* GridION runs from the NEMO dataset [11]. Test data comprised PromethION runs of the same three *E. coli* sample types and native and amplified DNA from *H. pylori* 26695. NEMO’s GATC and all-context models were evaluated on the same reads. More details are provided in Supplementary Methods.

## Data and code availability

All sequencing data used in this study are publicly available and listed in Supplementary Table S1. Code is available at https://github.com/guangzhaocs/Brieno, and model checkpoints are available at https://huggingface.co/guangzhaocs/Brieno.

## Acknowledgments and disclosure of funding

We acknowledge the generous computational resources provided by CSC – IT Center for Science, Finland. We thank the Research Council of Finland for funding (grant number 372081 to E.P.).

## Supplementary Information

**Supplementary Fig. S1:**
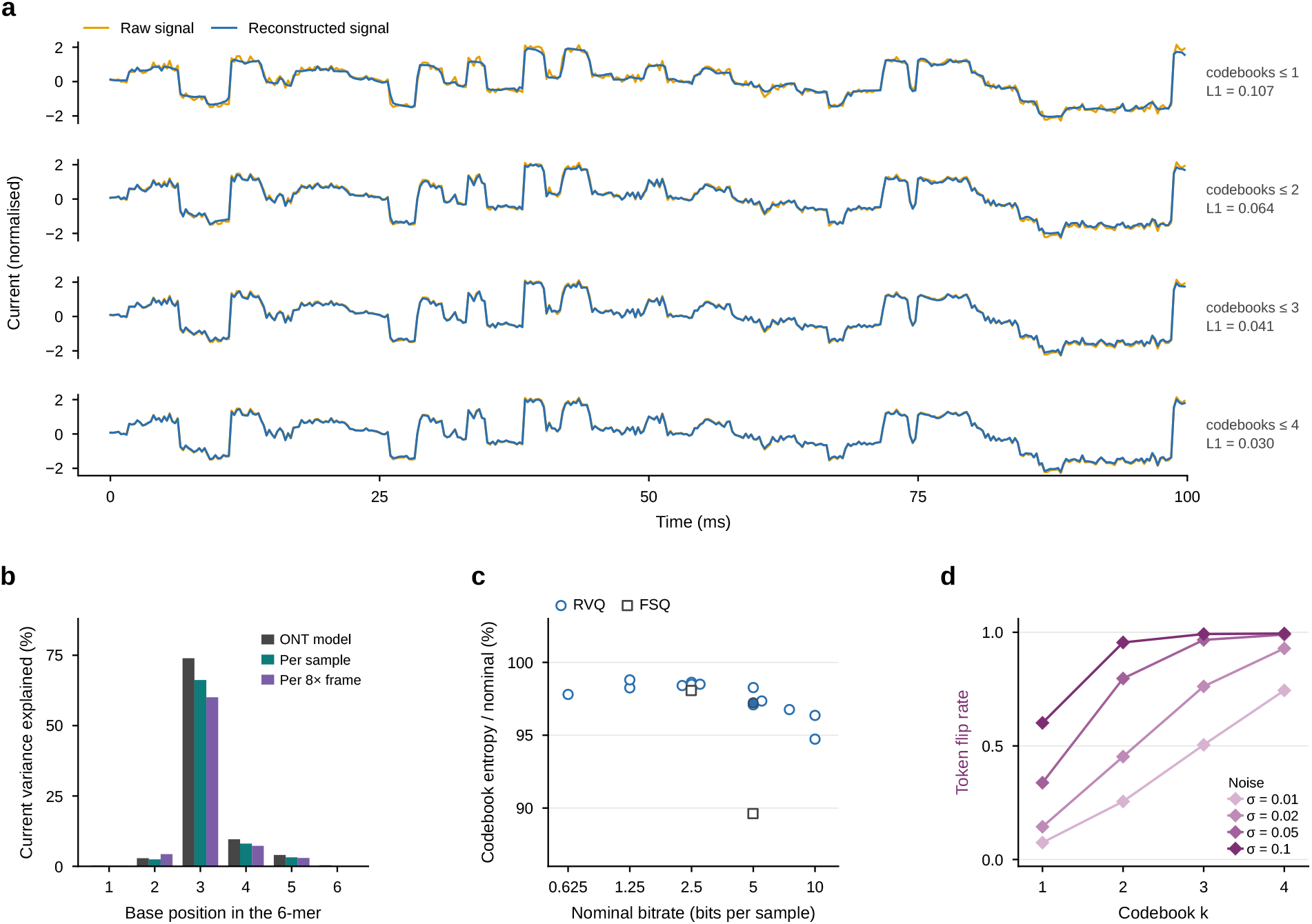
Characterization of the Brieno tokenizer. **a**, Comparison of raw (orange) and reconstructed (blue) signals using the first one, two, three or all four residual codebooks for the window shown in Fig. 2a. Reconstruction errors (*L*_1_) are indicated beside each trace. **b**, Percentage of current variance explained by each base position in the 6-mer, the fraction of the variance of the current accounted for by the identity of that base alone. Three independent routes are shown: the tabulated levels of the ONT pore model (4,096 6-mers), 6.5 million individual aligned signal samples from held-out lambda reads, and the 0.82 million 8 *×*downsampled frames those samples pool into. All three peak at the third position (73.9%, 66.2% and 60.1% respectively), which is therefore the base used to label reference positions throughout; the frame-level value remaining high indicates that 8 *×*pooling does not break the correspondence between one frame and one base. **c**, Codebook entropy as a percentage of the nominal bit rate, versus nominal bit rate, for residual vector quantization (RVQ; circles) and finite scalar quantization (FSQ; squares). The codebook entropy is the realized bit rate 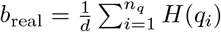, with 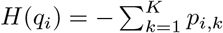 log_2_ *p*_*i,k*_ and *p*_*i,k*_ the frequency of entry *k* in codebook *i* over the validation set. The filled circle indicates the selected tokenizer, which realized 4.86 of its nominal 5.00 bits per sample. All RVQ tokenizers fell between 94.7% and 98.8%, so the nominal rate is a faithful upper bound on the realized one. Even at 10 bits per sample every codebook of the eight-codebook tokenizer carried 9.54–9.69 of its 10 bits, so the saturation in Fig. 2c does not reflect unused capacity. **d**, Fraction of tokens that change in each codebook after adding noise to the input signal at different standard deviations (*σ*).

**Supplementary Fig. S2:**
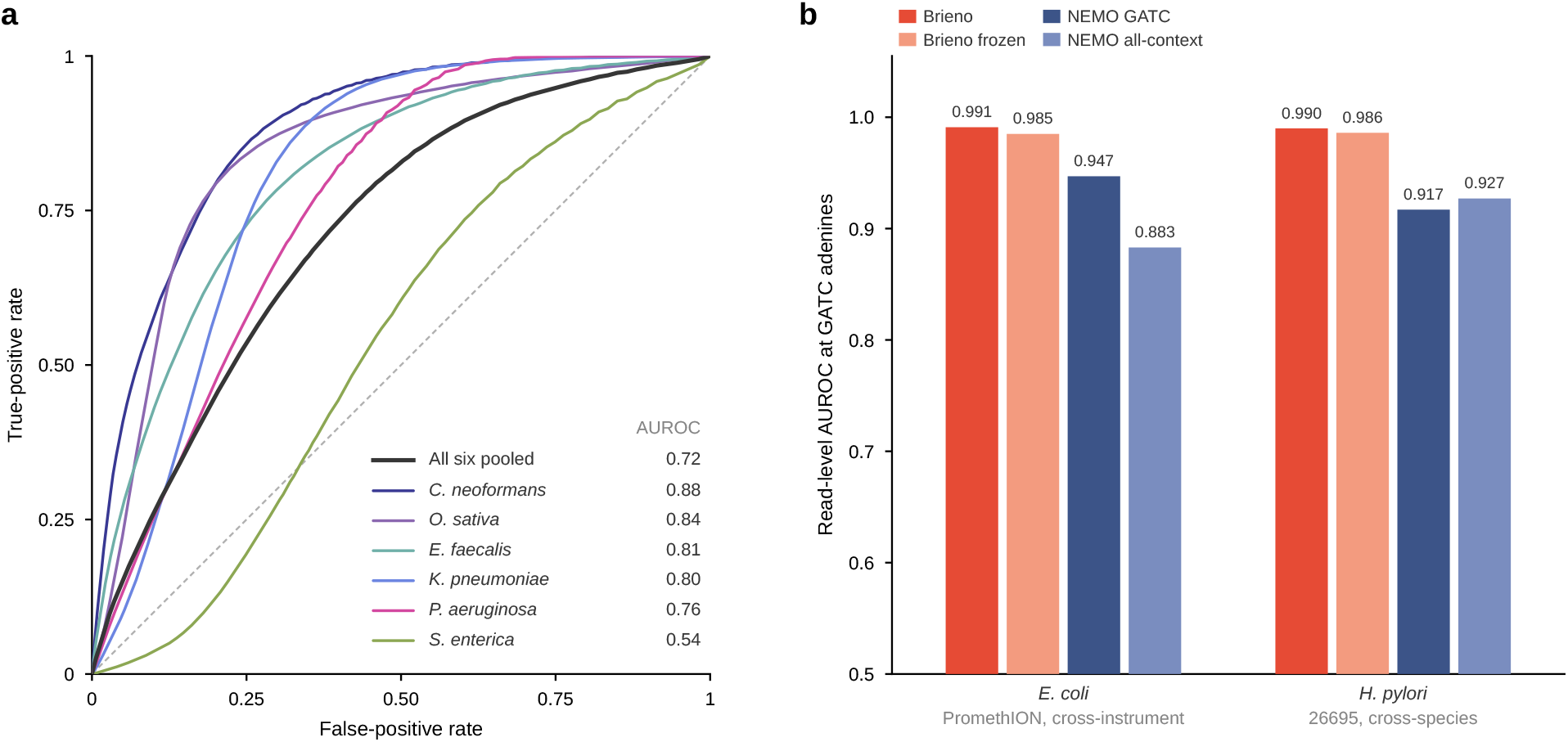
Detection of held-out species and 6mA detection at GATC sites. **a**, Receiver operating characteristic curves for flagging reads of species excluded from training of the twelve-class model. The novelty score is the negative margin between the two highest read-level mean log-probabilities. Positives are the 1,839,802 reads of the six held-out species, and negatives are the 3,390,616 reads of known species from sequencing runs unseen in training; each colored curve uses the reads of one held-out species against the same negatives, and the black curve pools all six. AUROC values are listed beside each curve, and the dashed diagonal indicates chance. Species colors match Fig. 3b. **b**, Read-level AUROC for 6mA detection at GATC adenines. Both Brieno models were trained on *E. coli* GridION runs comprising wild-type, *dam*^*−*^*dcm*^*−*^ mutant and whole-genome-amplified DNA, and evaluated on the same three sample types sequenced on a PromethION (six runs, cross-instrument) and on native versus whole-genome-amplified *H. pylori* 26695 (seven runs, cross-species). The cross-species evaluation is restricted to GATC adenines, the only sequence context that the NEMO GATC model calls. Both NEMO models [11] were applied to the same reads. The vertical axis starts at 0.5, the AUROC expected by chance. The NEMO all-context model was trained on *H. pylori*, so its value for that species is an in-distribution result rather than a cross-species one.

**Supplementary Fig. S3:**
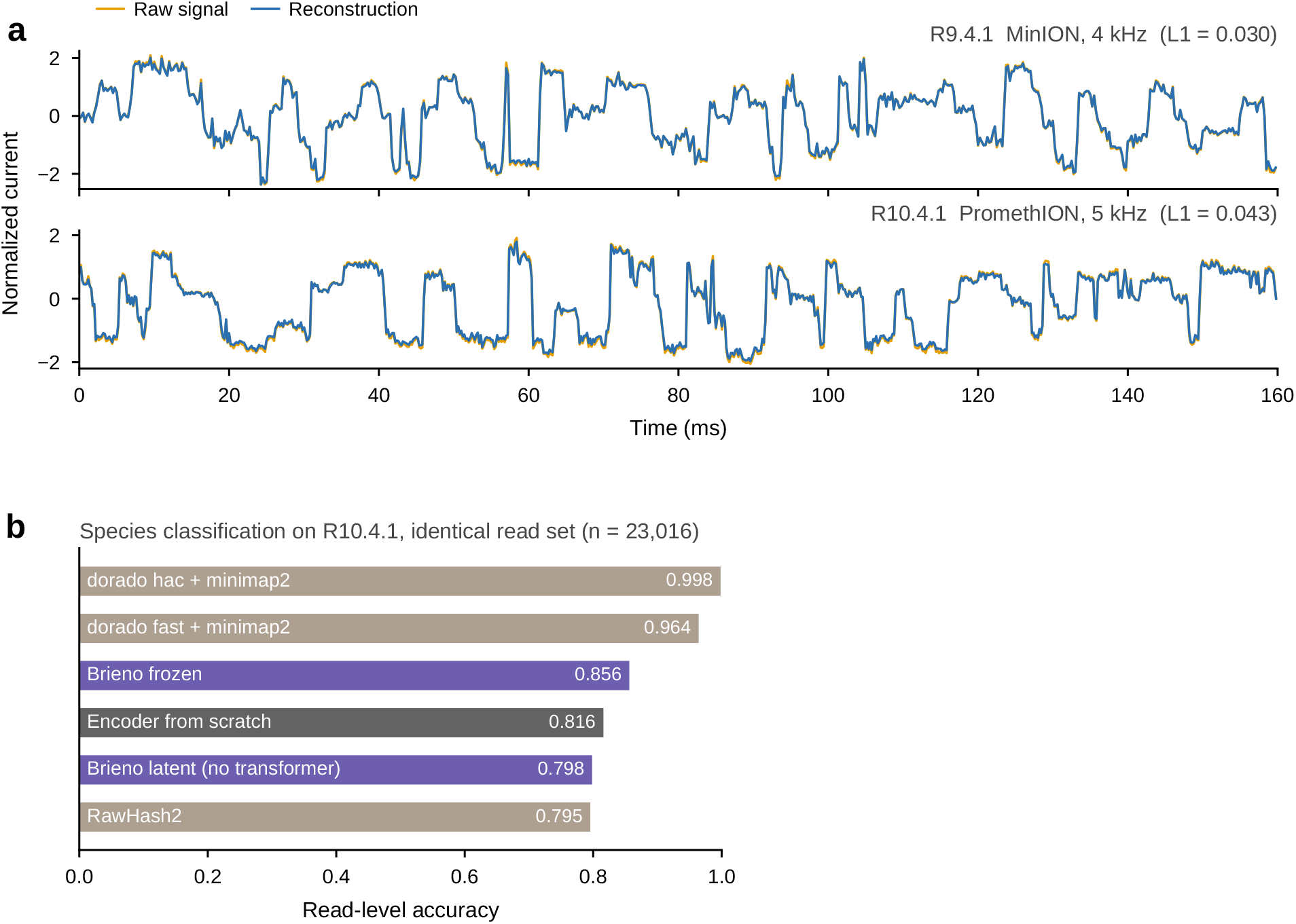
Signal reconstruction and species classification on R10.4.1 data. We used a public R10.4.1 flow cell of the ZymoBIOMICS HMW DNA standard (D6322), sequenced at 5 kHz using SQK-LSK114 (ENA ERR11924124, PromethION). Per-read species labels were obtained by aligning Dorado super-accuracy basecalls to the ten-species pan-reference using minimap2. **a**, Example 160 ms windows of raw signal (orange) and reconstruction (blue) from R9.4.1 and R10.4.1 data, using the tokenizer pretrained on R9.4.1 without further adaptation. Each trace is shown at its original sampling rate; *L*_1_ denotes the mean absolute reconstruction error over the displayed window. **b**, Read-level species classification accuracy after supervised adaptation to R10.4.1, with training and testing on disjoint reads from the same flow cell. All methods were evaluated on the same test reads (*n* = 23,016), with unmapped or unclassified reads counted as errors. Brieno with a frozen Transformer achieved an accuracy of 0.856, compared with 0.816 for the same encoder architecture trained from scratch and 0.795 for RawHash2. Using Brieno’s pooled continuous encoder features without the Transformer yielded an accuracy of 0.798. Dorado fast and high-accuracy basecalling followed by minimap2 achieved 0.964 and 0.998, respectively. Dorado super accuracy was excluded because its basecalls were used to derive the species labels.

**Supplementary Table S1:** Sequencing datasets used in this study. Public Oxford Nanopore recordings, R9.4.1 except the R10.4.1 flow cell in the last row. The pretraining corpus is the 16 datasets marked •: twelve were used for training and four were held out for validation. A ∘ marks data used only for fine-tuning and evaluation. The role column lists every stage a dataset enters, so a dataset reused across stages appears once. Rows are datasets or projects rather than individual runs. P, PromethION; G, GridION; M, MinION.

| Species / standard | Sample | Accession | Platform | Kit | Modification | Stage | Role |
|---|---|---|---|---|---|---|---|
| *H. sapiens* | HG001, ultra-long | ONT human WGS | P | ULK001 | 5mC | ● | Training |
| *H. sapiens* | HG001, ultra-long | ONT human WGS | M, G | RAD004 | 5mC | ● | Training; basecalling; 5mC |
| *A. thaliana* | Col-0 | PRJNA764549 | P | LSK109 | 5mC | ● | Training |
| *A. thaliana* | EcoGII/Hia5 | PRJEB69301 | M, P | LSK110 | 6mA | ● | Training |
| *Z. mays* | EcoGII | PRJEB69301 | P | LSK110 | 6mA | ● | Training |
| *O. sativa* | Nipponbare | PRJNA764549 | P | LSK109 | 5mC | ● | Training |
| *E. coli* | K-12 *dam*<sup>−</sup>*dcm*<sup>−</sup> | Zenodo 7970715 | M | LSK109 | none | ● | Training; basecalling |
| *S. aureus* | MRSA252 | PRJEB51164 | G | RBK004 | endogenous | ● | Training; basecalling |
| *Streptomyces* sp. S6 | – | PRJNA514949 | P | LSK109 | endogenous | ● | Training; basecalling |
| ZymoBIOMICS D6322 | HMW standard | PRJEB48692 | M | LSK110 | mixed | ● | Training |
| ZymoBIOMICS D6300 | mock standard | PRJEB29504 | G, P | LSK109 | mixed | ● | Training |
| Synthetic oligo | – | PRJEB76712 | G | LSK109 | 8-oxo-dG | ● | Training |
| *H. sapiens* | oxidised gDNA | PRJEB76712 | G | LSK109 | 8-oxo-dG | ● | Validation |
| *H. sapiens* | HG002, FAQ32498 | ONT gm24385 2021.09 | G | LSK110 | 5mC | ● | Validation |
| *E. coli* | CFT073 | PRJEB51164 | G | RBK004 | 6mA+5mC | ● | Validation |
| Lambda phage | VER5940 | PRJNA926802 | G | LSK109 | none | ● | Validation; basecalling test |
| ZymoBIOMICS D6300, D6322 | 7 runs, 3 laboratories | PRJEB22207, PRJEB29504, PRJNA604456 | M, G, P | LSK109 | mixed | ○ | Species identification |
| 18 species | 65 runs | 24 ENA/SRA projects | M, G, P | various | various | ○ | Species embedding |
| Enterobacteriaceae isolates | 8 flow cells, 188 isolates | PRJEB42440 | M, G | LSK109, NBD104/114 | native | ○ | Barcode demultiplexing |
| *E. coli* | 6 clinical isolates | PRJEB42440, PRJEB51164 | M, G | LSK109, RBK004 | native | ○ | Basecalling test |
| *K. pneumoniae* | 5 runs | PRJEB22207, PRJEB34353, PRJEB42440 | M, G | LSK108, LSK109, RBK004 | native | ○ | Basecalling test |
| *C. reinhardtii* | 5 runs | PRJEB31789, PRJEB59713 | M, P | LSK109, LSK110 | native | ○ | Basecalling test |
| *M. musculus* | ERR7222469 | PRJEB20569 | P | LSK110 | native | ○ | Basecalling test |
| *H. sapiens* | HG002, 3 flow cells | ONT gm24385 2020.11 | P | LSK109 | 5mC | ○ | Basecalling test; 5mC test |
| *H. sapiens* | HG002, FAP96083 and PAF27096 | ONT gm24385 2020.09 and 2021.09 | G, P | LSK109, LSK110 | 5mC | ○ | 5mC test |
| *E. coli*, *H. pylori* | 19 runs; WT, *dam*<sup>−</sup>*dcm*<sup>−</sup>, WGA | PRJNA1076669 | G, P | LSK109 | 6mA | ○ | 6mA training and test |
| ZymoBIOMICS D6322 | ERR11924124 | PRJEB65462 | P | LSK114 | mixed | ○ | R10.4.1 evaluation |

**Supplementary Table S2:**
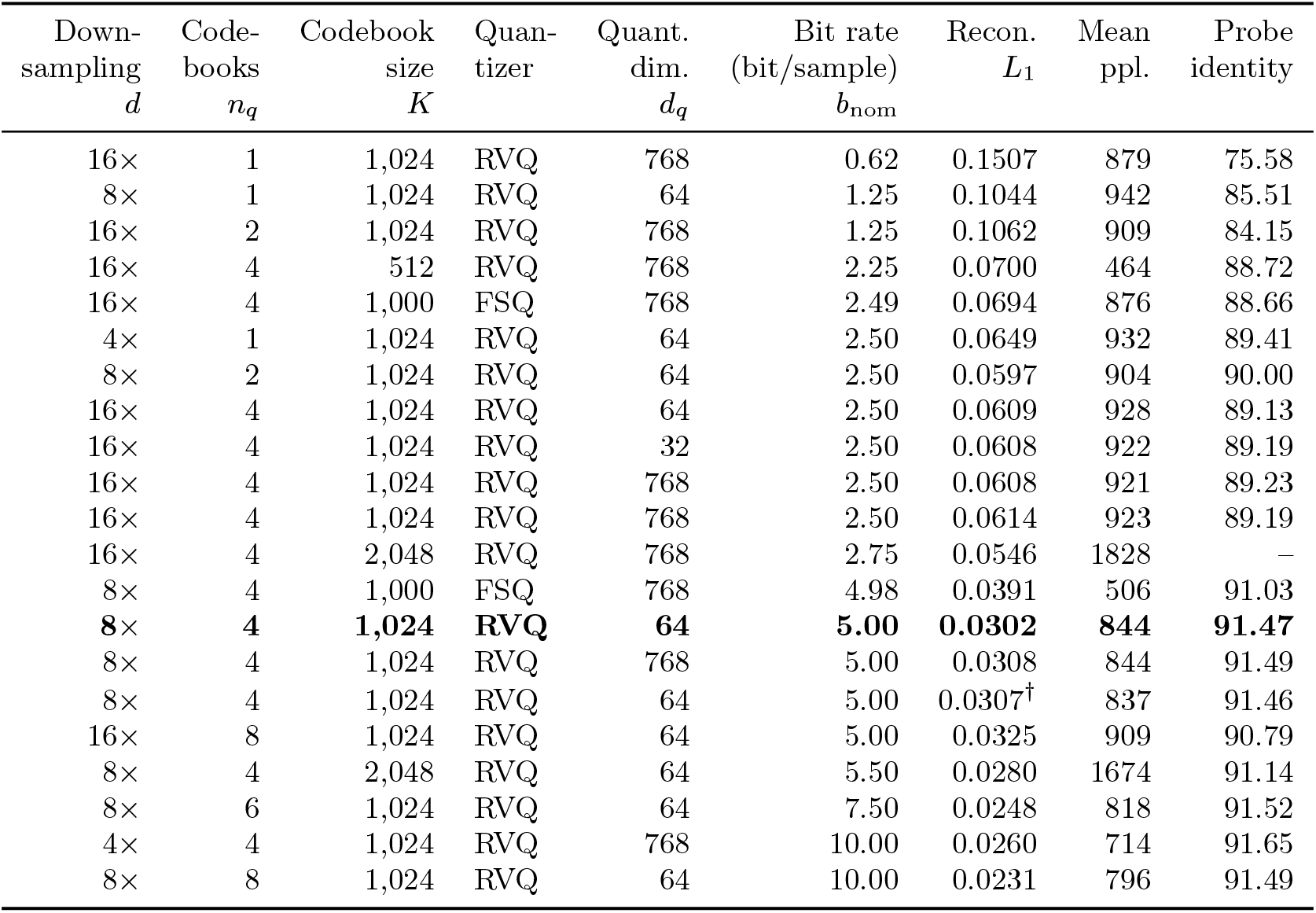
Configurations and performance of Brieno tokenizers. Comparison of 21 tokenizers with different downsampling factors, codebook numbers and sizes, quantization methods and quantization dimensions. Reconstruction error (*L*_1_) and mean codebook perplexity were evaluated on 405,502 held-out validation chunks at epoch 40. Basecalling probe identity (%) was evaluated on 20,000 held-out bacteriophage lambda chunks using frozen tokenizer features. Rows are ordered by nominal bit rate; the selected tokenizer is highlighted in bold. A dash indicates a probe that was not evaluated. Two tokenizer configurations were retrained with a second random seed to assess reproducibility: the two rows at 2.50 bits per sample differ only in seed and differ by about 1% in reconstruction error; the replicate of the selected tokenizer is not listed here. ^*†*^This run halves the encoder width to 384; every other tokenizer uses 768. RVQ, residual vector quantization; FSQ, finite scalar quantization.

**Supplementary Table S3:** Read-level basecalling accuracy. Matched bases divided by the length of the per-read reference sequence, the protocol used in the main text. Each dataset is scored on one read set shared by the four methods; *E. coli, K. pneumoniae* and *C. reinhardtii* are means over the runs listed.

| Dataset | Runs | Genome | Dorado fast | Dorado hac | Dorado sup | Brieno |
|---|---|---|---|---|---|---|
| Lambda | 1 | 48.5 kb | 0.9200 | 0.9432 | 0.9557 | **0.9546** |
| *E. coli* | 6 | 4.6 Mb | 0.9230 | 0.9383 | 0.9456 | **0.9446** |
| *K. pneumoniae* | 5 | 5.7 Mb | 0.9349 | 0.9520 | 0.9587 | **0.9538** |
| *C. reinhardtii* | 5 | 120 Mb | 0.9293 | 0.9492 | 0.9560 | **0.9508** |
| *M. musculus* | 1 | 2.76 Gb | 0.9113 | 0.9463 | 0.9572 | **0.9495** |
| HG002 PAG07162 | 1 | 3.1 Gb | 0.9453 | 0.9651 | 0.9722 | **0.9660** |
| HG002 PAG07165 | 1 | 3.1 Gb | 0.9410 | 0.9629 | 0.9700 | **0.9644** |
| HG002 PAG07506 | 1 | 3.1 Gb | 0.9165 | 0.9485 | 0.9590 | **0.9527** |

**Supplementary Table S4:**
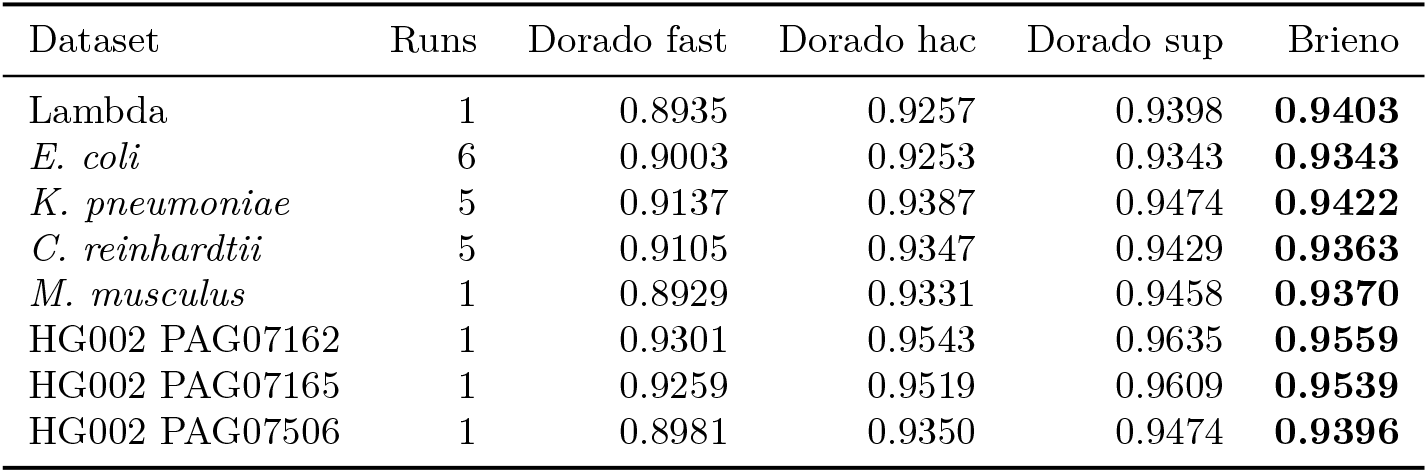
Read-level basecalling accuracy, alignment denominator. The same models, reads and runs as Supplementary Table S3, with matched bases divided by the alignment length instead of the reference length. Insertions and deletions therefore enter the denominator differently, which shifts every value.

| Dataset | Runs | Dorado fast | Dorado hac | Dorado sup | Brieno |
|---|---|---|---|---|---|
| Lambda | 1 | 0.8935 | 0.9257 | 0.9398 | **0.9403** |
| *E. coli* | 6 | 0.9003 | 0.9253 | 0.9343 | **0.9343** |
| *K. pneumoniae* | 5 | 0.9137 | 0.9387 | 0.9474 | **0.9422** |
| *C. reinhardtii* | 5 | 0.9105 | 0.9347 | 0.9429 | **0.9363** |
| *M. musculus* | 1 | 0.8929 | 0.9331 | 0.9458 | **0.9370** |
| HG002 PAG07162 | 1 | 0.9301 | 0.9543 | 0.9635 | **0.9559** |
| HG002 PAG07165 | 1 | 0.9259 | 0.9519 | 0.9609 | **0.9539** |
| HG002 PAG07506 | 1 | 0.8981 | 0.9350 | 0.9474 | **0.9396** |

**Supplementary Table S5:** Genome-level basecalling accuracy. The same models, reads and runs scored by the protocol of Wick et al. [2]: each read is aligned to the reference genome, a read that fails to map scores zero, and the denominator is the alignment length. The last column gives the unmapped fraction for each method.

| Dataset | Dorado fast | Dorado hac | Dorado sup | Brieno | Unmapped (%):<br>fast / hac / sup / Brieno |
|---|---|---|---|---|---|
| Lambda | 0.8737 | 0.9064 | 0.9399 | **0.9388** | 2.23 / 2.09 / 0.00 / 0.15 |
| *E. coli* | 0.8952 | 0.9238 | 0.9344 | **0.9313** | 0.58 / 0.16 / 0.00 / 0.31 |
| *K. pneumoniae* | 0.9102 | 0.9376 | 0.9474 | **0.9393** | 0.40 / 0.12 / 0.00 / 0.31 |
| *C. reinhardtii* | 0.9065 | 0.9319 | 0.9430 | **0.9333** | 0.45 / 0.30 / 0.00 / 0.31 |
| *M. musculus* | 0.7968 | 0.9309 | 0.9460 | **0.9318** | 11.24 / 0.25 / 0.00 / 0.56 |
| HG002 PAG07162 | 0.9279 | 0.9536 | 0.9635 | **0.9539** | 0.25 / 0.07 / 0.00 / 0.21 |
| HG002 PAG07165 | 0.9210 | 0.9510 | 0.9609 | **0.9513** | 0.60 / 0.09 / 0.00 / 0.26 |
| HG002 PAG07506 | 0.8543 | 0.9251 | 0.9475 | **0.9268** | 5.40 / 1.07 / 0.00 / 1.36 |

### Supplementary Methods

#### Tokenizer analysis

All tokenizers were trained for 40 epochs, each comprising 2.56 million chunks, using a batch size of 1,024, AdamW and a learning rate of 5 *×*10^*−*4^ with cosine decay. Fig. 2b and Supplementary Fig. S1 use 8,000 chunks from four species (lambda, *S. aureus, E. coli, Streptomyces*) aligned to their references at the signal level with Uncalled4 [7], giving 9.75 million reference positions and 16.1 million labeled frames. Each frame belongs to the reference position holding its center sample and is labeled with the third base of that position’s Uncalled4 6-mer, the position that explained the largest fraction of current variance (Supplementary Fig. S1b). The codebook entropy of the selected tokenizer reached approximately 97% of its nominal bit rate (Supplementary Fig. S1c), and later codebooks were more sensitive to input noise than the first, consistent with their role in encoding finer residual variation (Supplementary Fig. S1d).

Frame-level current is the mean of the eight samples in a frame, and the code-associated mean current of a codebook entry is the average frame-level current over all frames assigned that entry. Within-position differences were taken between the two frames of a reference position, restricted to positions covered by exactly two frames; between-position differences were taken between the mean currents of consecutive reference positions. Token flip rate was measured as the fraction of frames whose token changed after adding Gaussian noise of standard deviation *σ* to the normalized signal, over a 2,048-chunk subsample.

The basecalling probe that ranks tokenizers freezes the tokenizer and trains a small decoder on its tokens: a 256-dimensional codebook embedding, five bidirectional LSTM layers of 256 units and a CRF output, in fp32 for 60 epochs of 50,000 chunks. It was scored as the Needleman–Wunsch identity of each decoded chunk against its reference on 20,000 held-out lambda chunks.

#### Details of Transformer pretraining

Each prediction head is weight-tied to the embedding table of its own codebook: *p* is the softmax of 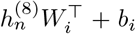 with *W*_*i*_ restricted to its first *K* rows, so the mask row is never a prediction target and only the bias *b*_*i*_ is a separate parameter. Training ran for 70 epochs, each comprising 10,000 optimization steps, on four GH200 GPUs, with a batch size of 80 chunks per GPU and two gradient-accumulation steps, yielding an effective batch size of 640. The optimizer is AdamW (*β* = (0.9, 0.98), *ε* = 10^*−*6^, weight decay 0.01) with a peak learning rate of 6 *×*10^*−*4^, 20,000 warm-up steps (two epochs) and cosine decay thereafter.

#### Details of species identification and barcode demultiplexing

The chunk head is a linear projection to 256 dimensions, three stride-2 convolutions (kernel 5, GroupNorm, GELU, dropout 0.1), the concatenation of an attention-pooled and a mean-pooled summary, and a two-layer MLP whose output layer has one unit per class, ten or twelve for species and 24 for barcodes. A read that yields *K* chunks *x*_1_, …, *x*_*K*_ receives one mean log-probability per class,

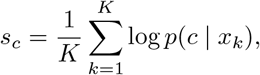

where *p*(*c*| *x*_*k*_) is the probability the head assigns to class *c* for chunk *x*_*k*_, and is assigned the class with the largest *s*_*c*_. Heads see a fixed number of chunks per class per epoch (50,000 for species, 12,000 for barcodes) with batch 128 and learning rate 5 *×*10^*−*4^ for 50 epochs, 30 for the fine-tuned Transformer, which converges faster, using AdamW, a one-cycle schedule and weight decay 0.05; the checkpoint with the best validation score is evaluated once on the test runs. The species training runs are two Birmingham GridION, two Birmingham PromethION and one Johns Hopkins MinION run. The per-read truth keeps reads with mean quality *≥*7, mapping quality *≥*20, identity*≥* 0.90, aligned fraction *≥*0.8 and at least one full chunk. UNCALLED [8], Sigmoni [14], Sigmap [15] and RawHash2 [9] were run with their published settings on the same reads and pan-reference, with unmapped reads counted as errors. The twelve-class model was trained with the same head on single-species and mock-community runs covering *A. thaliana, B. subtilis, C. reinhardtii, E. coli, H. sapiens, L. fermentum, L. monocytogenes, M. musculus, S. aureus, S. cerevisiae, Streptomyces* and *Z. mays*, with *C. neoformans, E. faecalis, K. pneumoniae, O. sativa, P. aeruginosa* and *S. enterica* held out; four of the six occur in mock-community runs that also carry training species, so their separation cannot be explained by run identity. Reads from the mock-community runs were labelled by species from the same Dorado super-accuracy and minimap2 alignment used for the ten-class benchmark. Pooled 512-dimensional read embeddings of 3,000 reads per species were projected with UMAP (30 neighbors, minimum distance 0.1, seed 0) for display only. For novelty detection the score is (*−s*_(1)_ *−s*_(2)_), the negative margin between the two largest values of *s*_*c*_, so that a small gap between the top two classes indicates a read the model cannot confidently place. The novelty AUROC was computed over held-out-species reads against the known-species reads of the same runs. For barcodes, the 24 native barcodes are EXP-NBD104 and EXP-NBD114 on SQK-LSK109 libraries, 5% of the training reads were held out as validation, and ground truth is the depositor’s sample assignment, each sample’s barcode recovered by majority vote of Dorado’s barcode calls (per-sample purity 94.8–99.0%). The input is the first 16,384 samples after adapter trimming, zero-padded for shorter reads. Dorado was run at fast, high and super accuracy, an unclassified read counting as an error, and the Deepbinner architecture was re-implemented and retrained on the same training flow cells with its original 1,024-sample window and scanning rule. For the coverage-matched comparison, test reads were ranked by the margin *s*_(1)_ *−s*_(2)_ between the two largest read-level mean log-probabilities and the least confident tail was discarded until the retained fraction equalled the fraction Dorado at super accuracy classified (0.9765); precision is the accuracy over the retained reads.

#### Details of basecalling

None of these test runs contributed training data; the human training data came from a different individual, HG001. The four training datasets are sampled with weights 0.40 (*E. coli*), 0.34 (HG001), 0.16 (*Streptomyces* sp. S6) and 0.10 (*S. aureus*). Training labels were produced by aligning Dorado super-accuracy calls to the reference with minimap2, keeping reads with mapping quality*≥* 50 and identity *≥*0.85, and realigning the signal with Uncalled4. The reported model was trained for 120 epochs of 50,000 chunks with batch 32, learning rate 10^*−*3^, 1,000 warm-up steps and cosine decay, in fp32 throughout. The representation comparisons of Fig. 4 use the same head at *h* = *d* = 256, trained for 60 epochs. Chunk-level identity is the Needleman–Wunsch identity of 20,000 held-out lambda chunks against their Uncalled4 labels. For read-level evaluation the per-read truth is cut from the genome using Dorado’s alignment with primary alignments only, mapping quality *≥*50, identity *≥*0.85 and aligned fraction *≥*0.8. Dorado v0.9.6 was run on the same reads using its fast, high-accuracy and super-accuracy models. In the genome-level evaluation, each basecalled read was instead aligned directly to the reference genome with minimap2, accuracy was calculated as matched bases divided by alignment length, and unmapped reads were scored zero.

#### Details of modification detection

The head is a linear projection to 256 dimensions, two convolutions (kernel 5) and a bidirectional LSTM of 192 units followed by a linear output, 1.55 M parameters. It emits one logit *g*_*n*_ per frame, and the logit of a base *b* whose Uncalled4 events span the frame set *ℱ*_*b*_ is

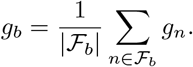

The loss is binary cross-entropy over every labeled base of the chunk, with non-CpG cytosines as negatives, and the scores of a base seen by overlapping chunks are averaged. The 5mC pseudo-labels keep CpG sites at which both ENCODE replicates are *≥* 95% or *≤*5% methylated with coverage *≥* 5; training ran for 16 epochs of 120,000 chunks with batch 32 and learning rate 5 *×*10^*−*4^. The EpiQC high-confidence sites are those at which EM-seq, TruSeq and SPLAT agree at *≥*90% or *≤* 10%. The five HG002 test flow cells are PAG07162, PAG07165 and PAG07506 (PromethION), FAP96083 (GridION, SQK-LSK110) and PAF27096 (PromethION); Dorado’s 5mCG model is the only R9.4.1 modification model released. For label efficiency a fixed, seeded fraction *f∈ {* 1, 0.1, 0.01} of every training run’s chunks was kept and each epoch sampled 120,000 chunks with replacement, so that the step budget was identical and only the number of distinct labeled chunks changed. The 6mA training data are the NEMO wild type, *dam*^*−*^*dcm*^*−*^ mutant and whole-genome-amplified *E. coli* runs, with GATC adenines labeled by sample type.

#### The representation ladder

Every downstream task feeds seven representations of the same chunk to the same head. A is the encoder output before quantization; B1 the sum of the four codebook vectors after back-projection; B2 the four token identities through a learned 256-dimensional embedding, initialized from the tokenizer’s codebooks; C1 the last-layer states of the frozen Transformer; C2 the same with the Transformer fine-tuned at learning rate 5 *×*10^*−*5^; D1 the same architecture as C2 with the Transformer randomly initialized instead of loaded, trained end to end with the head under the same data, schedule and optimization budget; and D2 the tokenizer’s encoder architecture with random initialization, trained end to end with the head. Tokenizer weights are frozen in A–D1, and D2 uses no tokenizer. Writing *s*(*·* ) for the test score of a rung, the differences reported in the text are

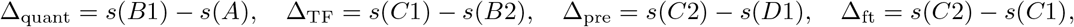

the cost of quantization, the contribution of the pretrained Transformer over its own tokens, the value of pretraining over random initialization and the gain from fine-tuning. On barcodes, D1 remained at chance for all 50 epochs with the training loss pinned near ln 24 = 3.178, and Transformer learning rates of 1 *×*10^*−*4^, 2 *×*10^*−*4^, 5 *×*10^*−*4^ and 1 *×*10^*−*3^ gave 0.0405, 0.0457, 0.0376 and 0.0374, while C1 and C2 reached 0.976 and 0.983 after a single epoch. Initializing the B2 embedding randomly rather than from the codebooks likewise left it at chance (0.0378), even at three times the training budget and four times the learning rate, and the same randomly initialized embedding stayed at chance when a bidirectional LSTM or a two-layer Transformer trained from scratch replaced the pooling head. Across the tested architectures and training settings the difficulty was specific to randomly initialized token embeddings and persisted as capacity, head architecture and the frozen or trainable state of the Transformer were varied; D2, which consumes the raw signal rather than token identities, trains normally on the same task.

## Notes

### Competing Interest Statement

The authors have declared no competing interest.

